# Cell-type-specific mapping of enhancers and target genes from single-cell multimodal data

**DOI:** 10.1101/2024.09.24.614814

**Authors:** Chang Su, Dongsoo Lee, Peng Jin, Jingfei Zhang

## Abstract

Mapping enhancers and target genes in disease-related cell types has provided critical insights into the functional mechanisms of genetic variants identified by genomewide association studies (GWAS). However, most existing analyses rely on bulk data or cultured cell lines, which may fail to identify cell-type-specific enhancers and target genes. Recently, single-cell multimodal data measuring both gene expression and chromatin accessibility within the same cells have enabled the inference of enhancer-gene pairs in a cell-type-specific and context-specific manner. However, this task is challenged by the data’s high sparsity, sequencing depth variation, and the computational burden of analyzing a large number of enhancer-gene pairs. To address these challenges, we propose scMultiMap, a statistical method that infers enhancer-gene association from sparse multimodal counts using a joint latent-variable model. It adjusts for technical confounding, permits fast moment-based estimation and provides analytically derived *p*-values. In systematic analyses of blood and brain data, scMultiMap shows appropriate type I error control, high statistical power with greater reproducibility across independent datasets and stronger consistency with orthogonal data modalities. Meanwhile, its computational cost is less than 1% of existing methods. When applied to single-cell multimodal data from postmortem brain samples from Alzheimer’s disease (AD) patients and controls, scMultiMap gave the highest heritability enrichment in microglia and revealed new insights into the regulatory mechanisms of AD GWAS variants in microglia.

## Introduction

The past two decades have seen significant advances in genome-wide association studies (GWAS), generating extensive catalogs of genetic variants linked to complex traits and diseases. However, over 90% of these identified variants are located in non-coding regions of the genome [1], and their disease-causing mechanisms remain largely unknown. Increasing evidence suggests that GWAS variants contribute to disease risk by modifying gene regulatory mechanisms in disease-relevant cell types [1, 2, 3]. Mapping enhancers, a principal class of gene regulatory elements, and its target genes has shown great promise in uncovering the functions of GWAS variants in specific cellular contexts [4]. However, most existing analyses utilize data from bulk tissues, which may fail to capture the highly cell-type-specific nature of enhancers, or data from cell lines, which may not accurately represent the biology of primary cell types and cells from diseased subjects [5, 6]. Some other analyses detect cell-type-specific enhancers using epigenetic data from a single modality, but they usually cannot identify the associated target genes due to the lack of data modality with gene expression measurements [5]. Recent technologies such as 3D epigenetic data and CRISPR screen data can be used to map enhancer-gene pairs in different cell types, but they remain laborious and costly to collect and may only be used to study cell lines [5, 7, 8].

The advent of single-cell multimodal technologies has unlocked unprecedented opportunities for mapping enhancers and target genes in specific cell types and contexts. Specifically, *paired* single cell assays for transposase-accessible chromatin using sequencing (scATAC-seq) and single cell RNA sequencing (scRNA-seq) allow for the profiling of both peak accessibility, a measurement of enhancer activity, and gene expression within the same cells. These data enable the identification of enhancer-gene pairs based on significant associations between peak accessibility and gene expression. As single-cell multimodal data are collected from both healthy individuals and those with various disease statuses across primary human cell types, it enables the computational inference of enhancer-gene pairs in a manner that is both cell-type-specific and context-dependent.

However, substantial computational and analytical challenges remain in the inference of peak-gene associations with single-cell multimodal data. From a computational perspective, the task is highly costly due to the extreme computational burden of screening all candidate peak-gene pairs in the genome. For example, using a cis-region of 1Mb to define candidate pairs [9] could result in *∼*10^4^ pairs to be tested in a single cell type. Meanwhile, most existing methods use Monte Carlo methods for statistical inference, such as sampling peaks from different chromosomes to construct null distributions [10] and using bootstrapping for uncertainty quantification [11]. For these methods, finding the *p*-value for a single peak-gene pair requires running the computational procedure *∼*10^2^ to *∼*10^4^ times. Combined with the large number of pairs and multiple cell types to consider, the overall computational costs of existing methods become prohibitively high, as the procedure needs to be run *∼*10^6^ to *∼*10^8^ times. From an analytical perspective, challenges arise due to the high sparsity, technical confounding and variations across biological samples in single-cell multimodal data. First, scATAC-seq data are highly sparse and often treated as binary [12], leading to the common practice of using binarized counts in peak-gene association inference [10, 11]. Recent evidence, however, suggests that peak counts contain quantitative information and count-based modeling can improve downstream analysis [12, 13]. As a result, the heuristic treatment of peak counts in existing methods may result in a loss of information and detection power in peak-gene association inference. Second, the confounding effects of varying sequencing depths pose a challenge for peak-gene association inference. It has been shown that varying sequencing depths can lead to spurious associations when using most existing methods for inferring gene-gene associations from scRNA-seq data [14]. For peak-gene associations, similar spurious associations may arise as both peak and gene counts correlate with their respective sequencing depths, and the sequencing depths from these two data modalities tend to be correlated (Supplementary Figure 1). Third, in the presence of multiple biological samples, inferring peak-gene associations are further challenged by coordinated variations in mean expression and accessibility across biological samples [15, 16, 17], leading to spurious associations. For example, consider a peak-gene pair that is not associated in sample A or sample B. If the mean expression and accessibility levels are both high in sample A and both low in sample B, then the peak-gene pair may appear associated if cells from samples A and B are pooled in the analysis without any careful adjustment.

To address these challenges, we present a new approach, called scMultiMap, that uses <u>s</u>ingle-<u>c</u>ell <u>multi</u>modal data to <u>map</u> cell-type-specific enhancer-gene pairs. scMultiMap is based on a multivariate latent-variable model that simultaneously models the gene counts and peak counts from single-cell multimodal data, and makes minimal parametric assumptions. It measures peak-gene association via the correlation between underlying gene expression and peak accessibility levels while accounting for variations in sequencing depths and across biological samples. Furthermore, we develop a highly computationally efficient moment-based estimation framework that provides both correlation estimates and theoretically justified analytical *p*-values for assessing statistical significance.

We evaluated scMultiMap by applying it to multiple paired scRNA-seq and scATAC-seq datasets, including datasets on peripheral blood mononuclear cells (PBMC) from healthy subjects and on postmortem brain samples from Alzheimer’s disease (AD) patients and controls. Our results show that scMultiMap maintains appropriate type I error control and achieves higher statistical power when compared with existing methods. Additionally, results from scMultiMap are more reproducible across independent single-cell multimodal data and also more consistent with results from orthogonal data modalities on the same cell type, such as promoter capture Hi-C [18], HiChIP [19] and proximity ligation-assisted chromatin immunoprecipitation sequencing (PLAC-seq) [3]. We demonstrated the superior computational scalability of scMultiMap by benchmarking its computing time on real data. To illustrate its utility in studying functions of GWAS variants in disease-related cell types, we applied scMultiMap to data collected on microglia from AD patients and controls. This analysis revealed high enrichment for AD heritability in microglia enhancers and identified enhancer-gene pairs containing selective AD GWAS variants [20], providing insights into the regulatory functions and disease-causing mechanisms of these AD variants in microglia.

## Results

### Overview of scMultiMap

We propose a joint latent-variable model to simultaneously model gene expression and peak accessibility. Suppose there are *p* genes, *q* peaks, and *n* cells in the cell type of interest. Let *x*_*ij*_ and *y*_*ij’*_ be the observed counts of gene *j* and peak *j*’ in cell *i*, respectively. Furthermore, let *z*_*ij*_ be the underlying expression level for gene *j*, defined to be the number of mRNA molecules from each gene relative to the total number of mRNA molecules in a cell. Let *v*_*ij’*_ be the underlying accessibility level for peak *j*’, defined to be the number of DNA fragments from each peak relative to the total number of DNA fragments in a cell. Use 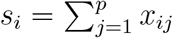 and 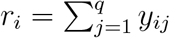 to denote the sequencing depths for scRNA-seq and scATAC-seq in cell *i*, respectively. We propose the following model

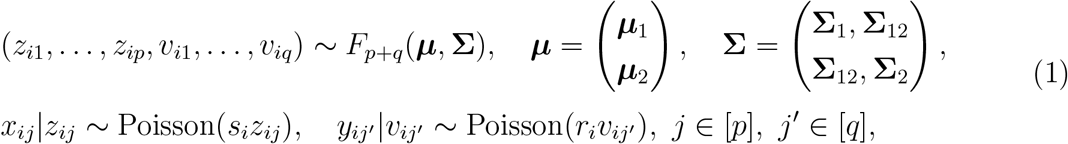

where *F*_*p*+*q*_(***µ*, Σ**) is a non-negative (*p* + *q*)-variate distribution with a mean vector ***µ*** of length *p* + *q* and a covariance matrix **Σ** of dimension (*p* + *q*) *×* (*p* + *q*). The covariance matrix **Σ** captures biological variations in the underlying gene expression and peak accessibility levels across cells, and is our main parameter of interest. Conditional on the latent expression level *z*_*ij*_ and accessibility level *v*_*ij’*_, gene count *x*_*ij*_ and peak count *y*_*ij’*_ are assumed to independently follow Poisson measurement models that depend on sequencing depths *s*_*i*_ and *r*_*i*_, respectively. Our approach does not impose specific parametric assumptions on *F*_*p*+*q*_(*·*),though(1) accommodates commonly considered distributions as special cases. For example, if *z*_*ij*_ (*v*_*ij’*_) follows a Gamma distribution, then *x*_*ij*_ (*y*_*ij’*_) follows a negative binomial distribution. In model (1), dispersion beyond the Poisson distribution is flexibly accommodated via *F*_*p*+*q*_(*·*).

Under model (1), we measure peak-gene correlations via **Σ**_12_, which directly quantifies the correlation strength between the underlying expression *z*_*i*_ = (*z*_*i*1_, …, *z*_*ip*_) and accessibility *v*_*i*_ = (*v*_*i*1_, …, *v*_*iq*_) and is not affected by variations in sequencing depths. When multiple biological samples (subjects) are present in the data set, we model the mean vector ***µ*** as ***µ***_*k*_ for subject *k*. This consideration accounts for variations in means across biological samples and avoids spurious associations (see Methods section).

We model peaks using a count measurement model that accommodates sequencing depth variations and overdispersion. Existing methods either treat peak data as binary [11] or free from sequencing depth variations [10], leading to a loss of power and potentially confounded estimates. Supported by recent evidence that Poisson modeling improves the downstream analysis of peak counts [12, 13] and the observation that peak counts display additional overdispersion compared to Poisson (Supplementary Figure 2), model (1) is able to better leverage the quantitative information in peak counts, remove potential confounding due to sequencing depths, and improve power in detecting association.

Estimation and testing under model (1) are non-trivial, as we do not wish to make restrictive parametric assumptions on *F*_*p*+*q*_(*·*). Moreover, there is usually a large number of peak-gene pairs to consider, making computational cost a practical and significant concern. To tackle these challenges, we propose a moment-based estimation framework that uses iteratively reweighted least squares (IRLS) with carefully specified weights to improve statistical efficiency. Under the proposed framework, we formulate a statistical test of hypothesis to evaluate the dependence between the underlying gene expression and peak accessibility levels from a given peak-gene pair, and analytically derive the null distribution. Correspondingly, the proposed test is theoretically justified and *p*-values can be analytically evaluated, without the need for time-consuming sampling-based inference. More details of the estimation and testing procedures can be found in the Methods section.

In summary, scMultiMap takes observed gene and peak counts as well as sequencing depths as input and generates analytical *p*-values for assessing the statistical significance of peak-gene associations. It properly models the distributions of gene and peak counts with a joint latent-variable model, accounting for variations in sequencing depths and across biological samples. The provided test has a controlled type I error rate, and enjoys a better statistical power. The computation of correlation estimates and *p*-values are fast and can be quickly implemented for tens of thousands of peak-gene pairs.

### scMultiMap has better detection accuracy and computational efficiency

To evaluate the performance of scMultiMap, we benchmarked its association detection accuracy and computational efficiency against existing methods using multiple single-cell multimodal datasets on PBMC from 10x Genomics [21, 22, 23]. In the benchmark analysis, we considered two peak-gene association inference methods that provide *p*-values, including Signac [10, 9] and SCENT [11] (Methods). In particular, Signac estimates peak-gene associations using Pearson’s correlations of gene counts normalized via sctransform [24] and raw peak counts. For testing, it constructs a null distribution by randomly sampling peak counts from other chromosomes. This procedure does not account for sequencing depth variations in peak counts, and has a computational cost of running *∼*10^2^ procedures for a single pair. Additionally, correlations of marginally normalized data are known to be biased by mean and overdispersion [14, 25], and random sampling is inadequate to adjust for such bias and may generate invalid *p*-values for inference. For SCENT, it employs a Poisson regression model that relates gene counts with binarized peak counts, and uses bootstrap methods for testing. This method may suffer from information loss due to the binarization of peak counts and confounding due to correlated sequencing depths. Additionally, it has a high computational cost ranging from *∼*10^2^ to *∼*10^4^ for a single pair [11].

To evaluate the type I error control of different methods, we construct null data by permuting peak counts randomly across cells. Correspondingly, the permuted peak counts are expected to be independent of gene counts (Methods). Figure 1A shows that scMultiMap has an appropriate type I error control with empirical type I errors matching the nominal level of 0.05. In comparison, the empirical type I errors of SCENT are slightly conservative for most pairs and notably inflated for some pairs. The empirical type I errors of Signac are substantially inflated for a subset of pairs. Next, to evaluate the detection power of different methods, we simulate representative single-cell multimodal data using model parameters estimated from PBMC data (Methods). The precision-recall curves in Figure 1B show that scMultiMap has higher power and a greater area under the curve. Both Signac and SCENT show reduced power, possibly because they are unable to fully extract the quantitative information in peak counts. Finally, to compare computational costs, we run all three methods on 729 CD14 monocytes and 31,132 peak-gene pairs using data from [26]. These pairs are derived from top 2,000 highly expressed genes, top 20,000 highly accessible peaks, and a cis-region of width 1Mb [9]. Figure 1C shows that the computational costs vary significantly across methods, with SCENT taking 1.37 days, Signac taking 1.56 hours, and scMultiMap taking only 12.77 seconds.

**Figure 1:**
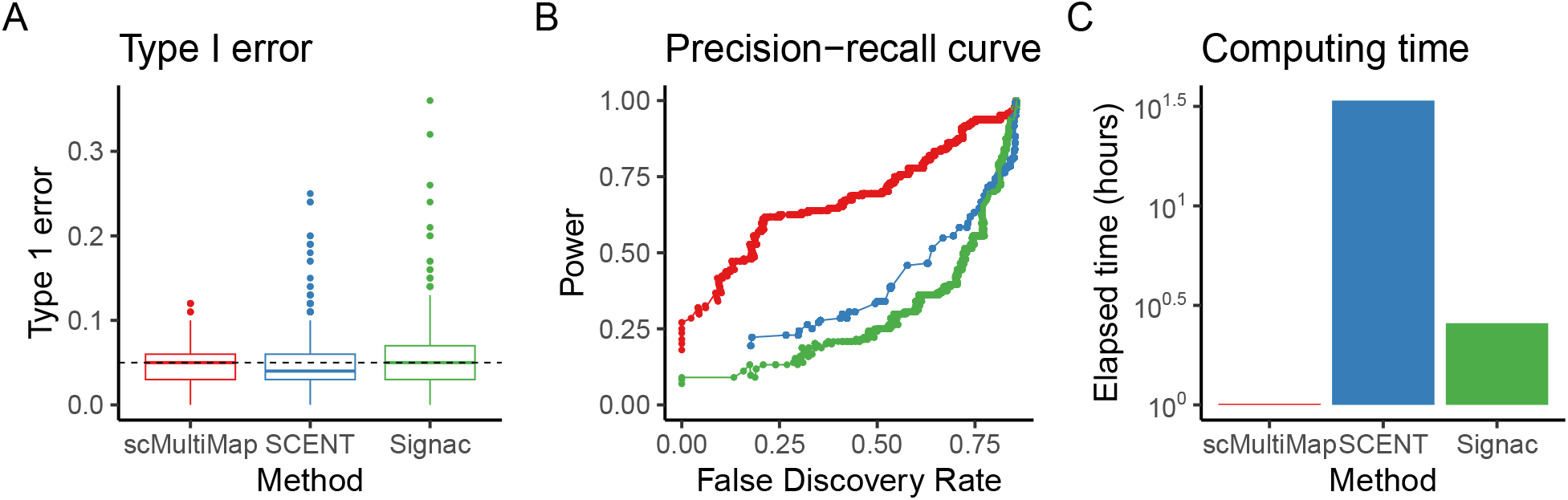
Performance evaluation of scMultiMap, SCENT and Signac using single-cell multimodal data on PBMC from 10x Genomics [21, 22, 23, 26]. A. Empirical type I errors on null data with independent gene expression and peak accessibility levels. The dashed line marks the nominal level of 0.05. B. Precision-recall curves on simulated data, with the same color legend as in A. C. Computing time in hours (log scale) on a dataset with 729 cells and 31,132 candidate peak-gene pairs.

### scMultiMap has higher reproducibility across independent datasets

The peak-gene association detection accuracy and power of scMultiMap are further evaluated through reproducibility analysis across multiple single-cell multimodal PBMC datasets. In particular, we evaluated reproducibility between biological replicates (different samples sequenced with the same instrument) and technical replicates (same samples sequenced with different instruments) [21, 22, 23] (Methods). Figure 2A shows that scMultiMap consistently yielded higher numbers of reproduced pairs in both reproducibility analyses across different BH-adjusted *p*-value cutoffs.

**Figure 2:**
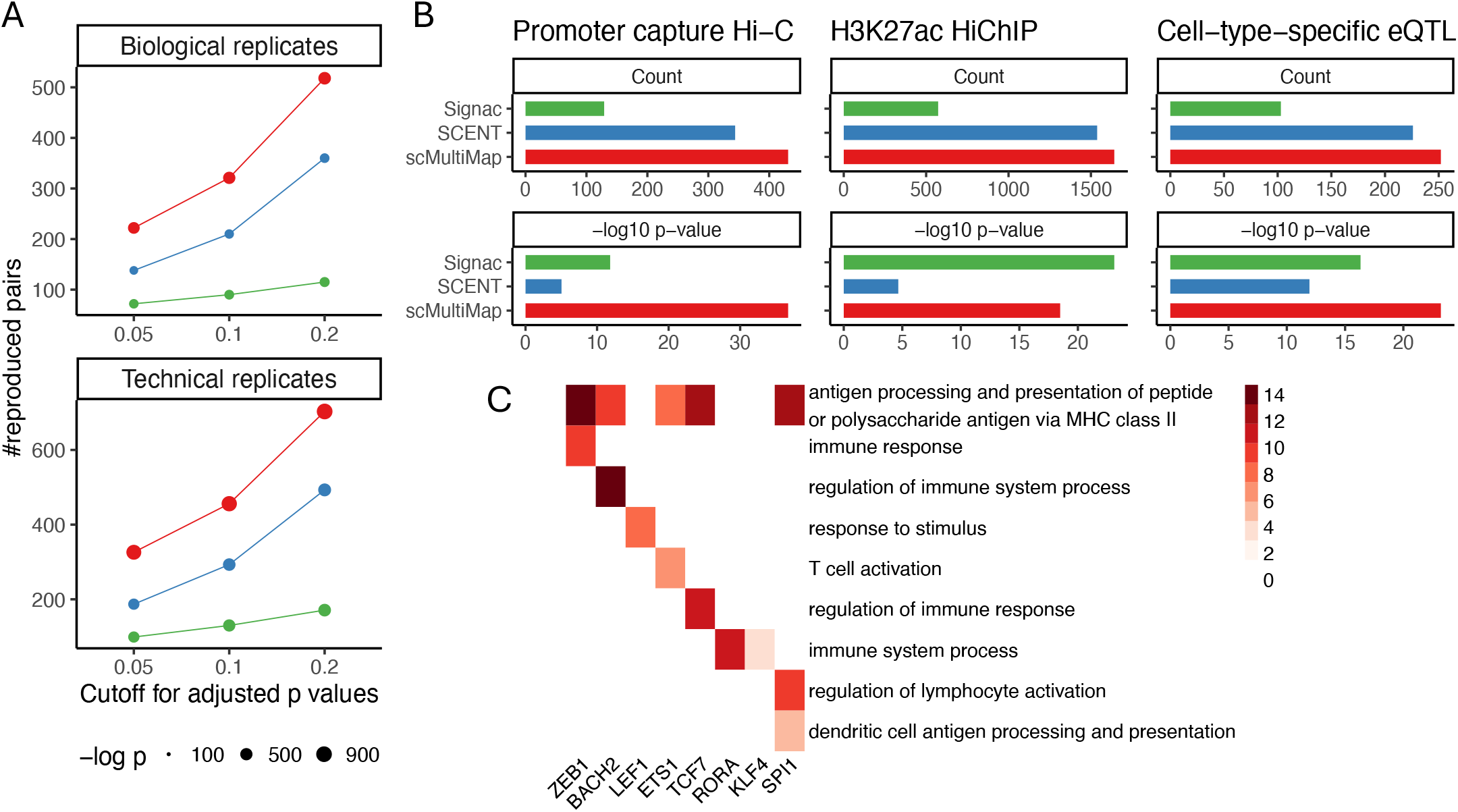
Comparison of reproducibility across methods and validation of regulatory trios inferred by scMultiMap. A. Number of significant peak-gene pairs reproduced between biological replicates and between technical replicates of single-cell multimodal data on CD14 monocytes across varying BH-adjusted *p*-values cutoffs. See B for the color legend. **B**. Consistency of scMultiMap findings with orthogonal data modalities. Significant peak-genes pairs on CD14 monocytes (BH-adjusted *p*-value *<* 0.1) were compared against enhancer-gene pairs measured by promoter capture Hi-C [18], H3K27ac HiChIP [19] and cell-type-specific eQTLs [27] (Methods). In A-B, statistical significance of the overlapped counts is evaluated with one-sided Fisher exact tests (−log *p* values are shown). C. Enrichment of GO biological processes among the target genes in trios identified for each TF in CD14 monocytes.

Furthermore, we compared the enhancer-gene pairs inferred from multimodal datasets with those inferred from orthogonal data types on the same cell type, including promoter capture Hi-C data, HiChIP data and cell-type-specific eQTL data (Methods). Figure 2B shows that peak-gene pairs identified by scMultiMap are the most consistent with promoter capture Hi-C [18], H3K27ac HiChIP [19], and cell-type-specific eQTL [27] data on CD14 monocytes. These pairs show the largest numbers of overlaps, and the overlaps are statistically significant (Methods).

Finally, we illustrate the utility of scMultiMap in studying gene regulation in specific cell types. For this task, we constructed gene regulatory trios [28], consisting of transcription factors (TFs), peaks and target genes, where a TF binds to the motif within the peak to regulate a target gene in close proximity to the peak. We quantified the association between TF and peak, peak and target gene with scMultiMap, and the association between TF and target gene with CS-CORE [14]. We considered the trios with simultaneous significant associations on TF-peak, peak-gene and TF-gene as cell-type-specific gene regulatory trios (Methods). Figure 2C shows the Gene Ontology (GO) enrichment in biological processes among the target genes in identified trios for the eight top enriched TFs in CD14 monocytes. Consistent with the role of monocytes as antigen-presenting cells [29], the pathway for antigen processing and presentation via MHC class II was found to be strongly enriched, regulated by multiple TFs. Many other immune functions of CD14 monocytes were also identified, such as T cell and lymphocyte activation [30, 31].

### scMultiMap identified biologically relevant gene regulatory mechanisms in brain cells

We further demonstrate the utility of scMultiMap with an analysis of the single-cell multimodal data collected from postmortem brain samples of Alzheimer’s disease patients and controls [32]. We focus on the cells from healthy controls. This part of the data include cells from eight subjects (biological samples), introducing potential variations across subjects and representing a more challenging scenario when compared to the PBMC data, which involves only one subject.

Using a bootstrap-based analysis (Supplementary Methods), we found significant variations in mean gene expression and mean peak accessibility across subjects (Supplementary Figure 3). Interestingly, these variations tend to be correlated between peaks and genes (Supplementary Figure 4), leading to spurious associations if left unaddressed. To understand the impact of coordinated variations across subjects on existing methods, we permuted real data to generate a null setting with independent peak accessibility and gene expression while preserving across-subject variations, and then evaluated the empirical type I errors (Methods). Figure 3A shows that Signac now exhibits much higher inflation in type I errors than in the PBMC data, as its correlation metric fails to account for variations across samples. SCENT also shows inflated type I errors, despite including subject id as a covariate in its regression model. In contrast, the empirical type I errors of scMultiMap align with the nominal level. We also evaluated the power of scMultiMap on simulated data with variations across subjects (Methods). Figure 3B shows that scMultiMap maintains the highest power. Notably, there is a larger gap between SCENT and Signac compared to the PBMC data (Figure 1B), as Signac is more confounded by the coordinated variations between peaks and genes across subjects.

**Figure 3:**
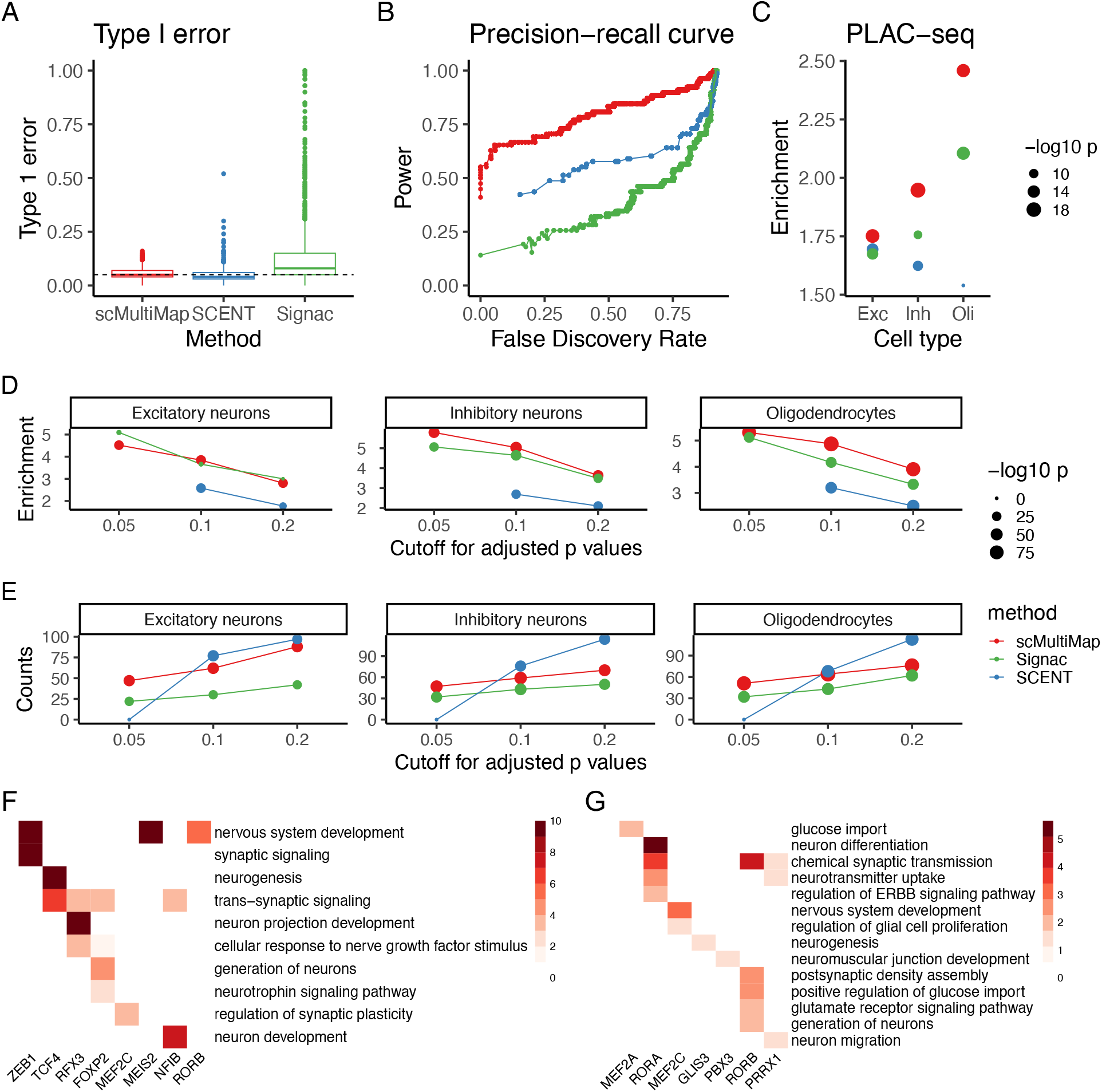
Results from scMultiMap on single-cell multimodal data from postmortem brain samples in [32]. A. Empirical type I errors on permuted data. Normalized peak values were randomly permuted within subject to break dependency with gene expressions while preserving variations among subjects. The dashed line marks the nominal level of 0.05. B. Precision-recall curves on simulated data. See color legend in A. C. Consistency of significant pairs (BH-adjusted *p*-value *<* 0.2) with enhancer-gene pairs measured by PLAC-seq [3] in excitatory neurons (Exc), inhibitory neurons (Inh) and oligodendrocytes (Oli) (Methods). See color legend in A. D-E. Reproducibility of significant pairs with independent singlecell multimodal data on brain samples from [33] across cutoffs of BH-adjusted *p*-values, as evaluated by the enrichment (D) and the number (**E**) of reproduced counts. In C-E, enrichment is quantified by odds ratio (OR) and log OR in C and D respectively (OR=0 not shown), and *p*-values are from one-sided Fisher exact tests. F**-G**. Enrichment of GO biological processes among the target genes in trios identified for each TF in excitatory neurons (**F**) and astrocytes (G). Color intensity is given by BH-adjusted -log_10_ *p*-values from 11 one-sided Fisher exact tests (values larger than 10 were set to 10).

We then evaluated the accuracy of scMultiMap on brain data through consistency and reproducibility analysis with other datasets measuring enhancers and target genes in brain cell types. Due to the large inflation (Figure 3A) and systematic bias in Signac correlations by mean levels [14, 25], we applied a permutation-based procedure to Signac to ensure a fair comparison with other methods (Methods). For consistency analysis, we used PLACseq data from [3], which measures the interaction between promoter and distal regulatory regions in neuronal cells, oligodendrocytes and microglia using cell nuclei isolated from human brains. We compared the significant peak-gene pairs identified in single-cell multimodal data with PLAC-seq results (Methods). Figure 3C shows that scMultiMap yields the highest enrichment for consistent enhancer-gene pairs in the three most abundant brain cell types, excitatory neurons, inhibitory neurons, and oligodendrocytes. The numbers of consistent pairs in microglia are insignificant for all methods (raw *p*-value *>* 0.05), possibly due to the smaller number of microglia in the single-cell multimodal dataset.

We further analyzed a second single-cell multimodal dataset from postmortem brain samples [33] and evaluated the reproducibility of findings with those from data in [32] (Methods). We focused on the three abundant brain cell types as in Figure 3C and did not include astrocytes and microglia due to the limited number of cells from these two cell types in the second dataset [33]. Figures 3D-E show that scMultiMap generally has the highest enrichment and the largest number of overlapping pairs between the two independent datasets. While Signac has similar enrichment as scMultiMap, it has lower power and identified fewer pairs compared to scMultiMap. SCENT failed to generate reproducible discoveries when the significance cutoff is stringent, and the enrichment of reproduced discoveries is much lower when the significance cutoff is lenient. This suggests that SCENT may suffer from lower power in identifying true pairs and inflated false positive discoveries that cannot be replicated across datasets.

We further applied scMultiMap to identify regulatory trios and studied the biological processes regulated by enriched TFs in five brain cell types (Figures 3F-G, Supplementary Figure 5) (Methods). The identified enriched GO biological processes are consistent with existing literature on the functions of these TFs in excitatory neurons (Figures 3F). For example, FOXP2 is known for its regulatory role in neurite outgrowth [34] and neuronal differentiation [35], MEF2C regulates synapse number and function to facilitate learning and memory [36], and RORB contributes to the establishment of neocortical layers [37]. Similarly, the trios yield biologically plausible enrichment in astrocytes (Figures 3G). For instance, chemical synaptic transmission, a key function of astrocytes [38, 39], is found to be enriched among the target genes of three TFs in Figure 3G. Strong enrichment for multiple biological functions was identified for RORA, an important pluripotent transcription factor that supports the neuro-protective and anti-inflammatory role of astrocytes in the brain [40]. The trios also yielded enrichment for cell-type-specific functions in oligodendrocytes (e.g. axon ensheathment, lipoprotein metabolic process), inhibitory neurons (e.g. chemical synaptic transmission, migration of Purkinje cell) and microglia (e.g. regulation of gliogenesis, inflammatory response) (Supplementary Figure 5).

### scMultiMap mapped GWAS variants of Alzheimer’s disease to target genes in microglia

Previous AD GWAS have identified 75 loci associated with the risk of developing AD [41]. However, the functional pathways and the cellular contexts in which these variants exert their effects remain unclear. Here, we leverage the candidate enhancer-gene pairs identified from scMultiMap in cells from healthy control and AD subjects to study cell-type and context-specific target genes of AD GWAS variants. We focus on microglia, the innate immune cell types in the brain whose cis-regulatory elements are the most enriched for AD genetic risk [3, 42].

First, we demonstrate the power of scMultiMap in identifying candidate cis-regulatory elements in microglia via AD heritability enrichment analysis. Given the high microglia-specific enrichment reported in the literature [3, 42], we hypothesize that a more powerful and accurate method for identifying cis-regulatory elements should yield higher heritability enrichment for AD in microglia. It is important to note that scMultiMap can capture general cis-regulatory elements that modulate the expression of target genes, even though it was initially motivated by detecting enhancer-gene pairs. Figure 4A shows that using peaks from significant peak-gene pairs, scMultiMap generated higher and more significant AD heritability enrichment in a stratified linkage disequilibrium score (S-LDSC) analysis based on three AD GWAS studies [41, 44, 45] (Methods).

**Figure 4:**
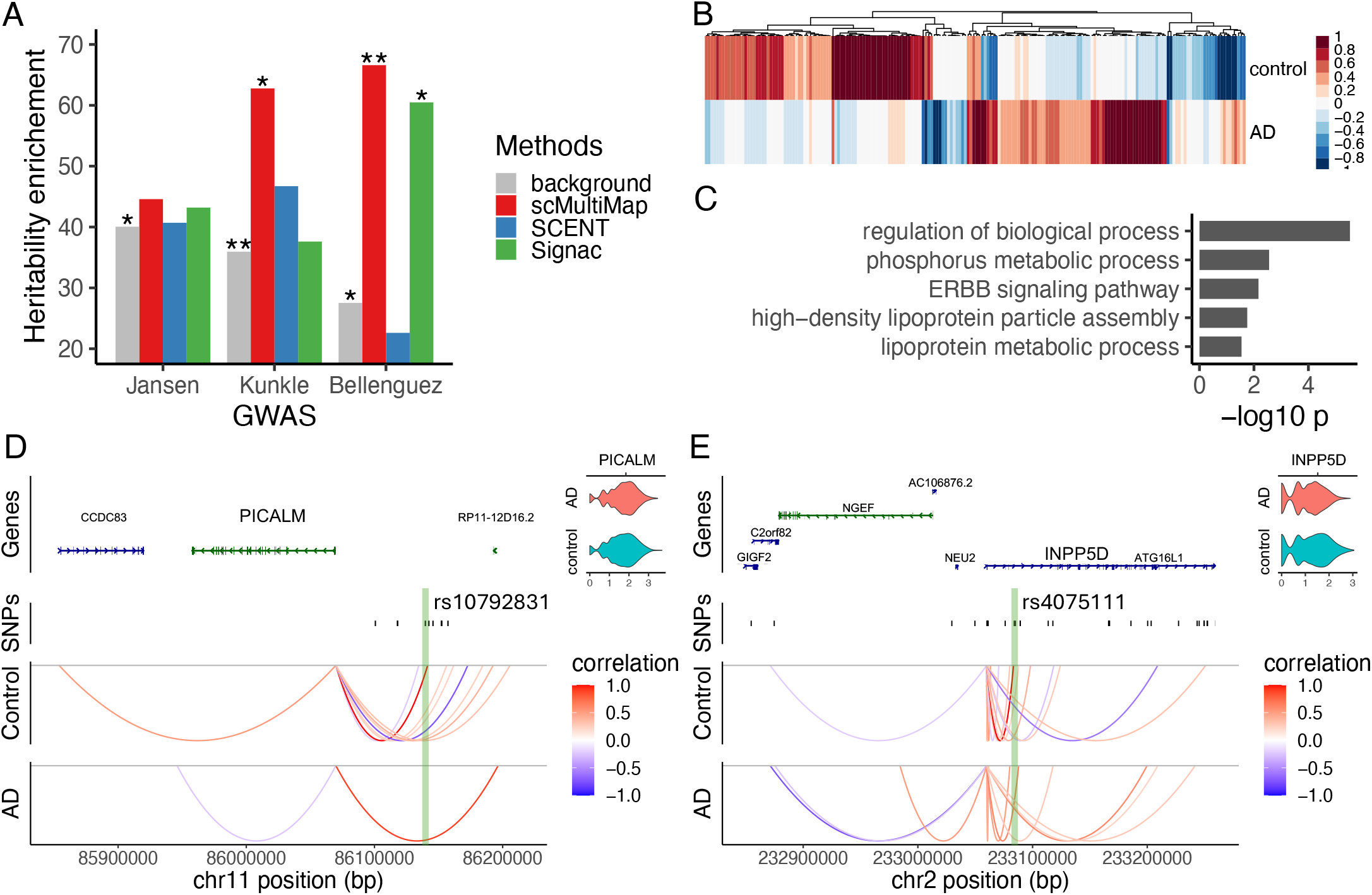
Studying the functional role of selective AD GWAS variants in microglia with scMultiMap. A. AD heritability enrichment for peaks from significant peak-gene pairs (raw *p*-value *<* 0.05) in microglia. Summary statistics from three AD GWAS studies were used: Jansen [44], Kunkle [45] and Bellenguez [41].* and ** denote *p*-value *<* 0.1 and 0.01 respectively for one-sided *p*-values of heritability enrichment from S-LDSC. B. Differential peak-gene pairs in microglia from control and AD subjects. C. Enrichment of GO biological processes among the genes from significantly differential peak-gene pairs. BH-adjusted *p*-values from one-sided Fisher exact tests are shown. D. scMultiMap mapped AD variant rs10792831 to PICALM in microglia from control subjects and the association is insignificant in microglia from AD subjects. E. scMultiMap mapped AD variant rs4075111 to INPP5D in microglia from control subjects and the association is insignificant in microglia from AD subjects.

We then compared the inferred candidate peak-gene pairs in microglia from healthy controls and AD subjects. We identified significantly differentially associated peak-gene pairs (Figure 4B) through a permutation analysis (Methods). We further intersected genes in significantly differentially associated peak-gene pairs (raw *p*-value *<* 0.05) and differentially expressed genes from [32] and evaluated the enriched GO biological processes. Figure 4C shows the top enriched processes, three of which are closely related to lipid metabolism—a pathway previously implicated in AD pathogenesis and microglia [44, 46, 47]. Similar analysis on astrocyte, another glial cell type critical to AD disease mechanisms [48, 49, 50], also revealed differentially associated peak-gene pairs and implicated pathways in lipid metabolism and neurofibrillary tangle assembly, both of which have been linked to AD in astrocytes [49, 50] (Supplementary Figure 6).

We next integrated the scMultiMap findings with AD GWAS variants to study the functional roles of these variants in microglia. We leveraged a set of AD GWAS variants that have been fine-mapped and prioritized based on microglia 3D epigenome annotations from a recent study [20]. scMultiMap found two significant peak-gene pairs that overlap with the selective GWAS variants and their putative target genes from [20] (Figures 4D-E). In particular, these pairs were validated by promoter capture Hi-C data on human pluripotent stem cell-derived microglia-like cells [20], and they were not found by Signac or SCENT (raw *p*-value *>* 0.05). Figure 4D shows that scMultiMap identified a peak containing the AD GWAS variant rs10792831 as associated with PICALM in microglia. PICALM is a known AD locus with upregulated expression in microglia from brain samples of AD patients [51]. The variant rs10792831 is located 72 kb away from the transcription start site of PICALM. According to the FAVOR database [52], this variant has a high epigenetics active score (with elevated H3K27ac and H3K4me1 levels from ENCODE [53]), a high TF score for overlap with TF binding sites, and a CADD score [54] of 10.11 for deleteriousness. While these results suggest moderate regulatory potential without reference to cellular contexts, scMultiMap highlights its functional role in microglia and identifies the target gene regulated in this cell type. Interestingly, this pair (the peak with rs10792831 and PICALM) is also among the significantly differentially associated peak-gene pairs in microglia, where the association is no longer significant when microglia from AD samples are considered (Figure 4D). This implies that the identified candidate enhancer is context-specific and that rs10792831 may contribute to AD risk by dysregulating the enhancer and, consequently, the expression of PICALM in microglia. Figure 4E shows that scMultiMap identified another peak containing the AD GWAS variant rs4075111 as associated with INPP5D in microglia. INPP5D is a known AD locus with downregulated expression in microglia from AD samples [32]. The associated variant, rs4075111, is located within the intronic region of INPP5D, 24 kb away from the transcription start site. It has been identified as being within the transcription factor binding site and enhancer for INPP5D by GeneHancer [55], and it has a high epigenetics active score based on FAVOR [52]. scMultiMap further highlights this enhancer-gene pair in the specific cellular context of microglia. Moreover, the association is not significant in microglia from AD patients, suggesting the context-specificity of the enhancer and the potential regulatory role of the AD GWAS variant rs4075111 in microglia.

## Discussion

We have introduced scMultiMap, a new statistical method for mapping cell-type-specific enhancer-gene pairs using single-cell multimodal data. By properly modeling peak counts and confounders in sequencing experiments, scMultiMap demonstrates higher statistical power to detect true enhancer-gene pairs and is robust against false positive associations due to varying sequencing depths and variations across biological samples. Utilizing a moment-based estimation framework and theoretically derived statistical tests, scMultiMap provides analytical *p*-values and has computational complexity that is less than 1% of existing methods. Systematic simulations and real data analyses show that scMultiMap better identifies reproducible and externally validated enhancer-gene pairs, making it a valuable tool for studying gene regulation in cell types. Integrative analysis with AD GWAS variants illustrates that scMultiMap can offer functional insights into the regulatory roles of GWAS variants in disease-related cell types, generating hypotheses for downstream validation and identifying potential targets for therapeutic intervention.

While scMultiMap is designed to quantify pairwise associations between peaks and genes, it can also serve a tool for constructing gene regulatory networks [56, 57, 58]. The gene regulatory trios illustrated here can be adapted to infer networks of target genes regulated by TFs. In this work, we have focused on paired scRNA-seq and scATAC-seq as an example of single-cell multimodal data, partly due to its availability. However, scMultiMap can also be applied to other multimodal sequencing data, such as those that jointly profile transcriptome and histone modification in single cells [59, 60], to study gene regulation in cell types.

We note that for the purpose of identifying enhancers, the results in this work are limited by the use of chromatin accessibility, which is necessary though not sufficient to define enhancers [3]. Additional data modalities, especially histone modification data in the cell types of interests, are needed to further validate the prioritized enhancers. As aforementioned, scMultiMap may also be applied to infer the association between gene expression and histone modification across single cells [59, 60], which will offer complementary insights into enhancer-gene pairs compared to those inferred from paired scRNA-seq and scATAC-seq data.

We have demonstrated the improved power of scMultiMap for identifying peak-gene pairs compared to existing methods. However, the power of this task is still limited by the sparsity and the small number of cells in single-cell multimodal data. It has been established that larger numbers of cells are needed for detecting over-dispersion [61], which is the variance of underlying abundance in the distribution *F* in (1). Similarly, for association inference, larger numbers of cells are also necessary to accurately estimate and test the covariance between underlying abundances. Often, there are fewer than a few hundred cells from the cell type of interest, providing limited information and yielding low to moderate power for association inference. Fortunately, larger consortium efforts are ongoing, collecting single-cell multimodal data with more cells across more biological samples. This will greatly improve the power of mapping enhancer-gene pairs in a cell-type-specific and context-specific manner. Given its better power and computational scalability, we expect scMultiMap to be a useful and practically appealing tool for analyzing these new consortium data, studying gene regulation, and elucidating the regulatory functions of GWAS variants in cell types.

## Method

### scMultiMap method

From model (1), it holds for the count of gene *j* that

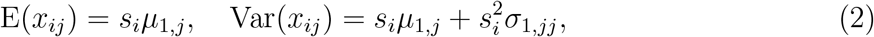

for the count of peak *j*’ that

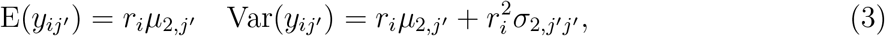

and for counts from gene *j* and peak *j*’ that

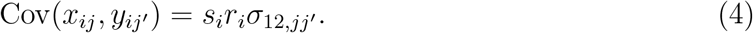

The estimation in scMultiMap includes two steps. The first step estimates the mean and variance parameters in (2)-(3) with IRLS. For example, for peak *j*’, we derive the following regression equations based on (3):

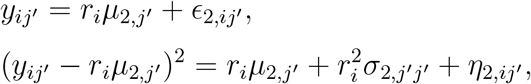

where mean-zero error terms *ϵ*_2,*ij’*_’s are independent across *i* for peak *j*’, and the same for *η*_2,*ij’*_’s. We propose to estimate *µ*_2,*j’*_ and *σ*_2,*j’j’*_ with weighted least squares estimators 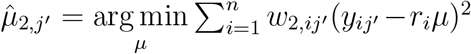 and 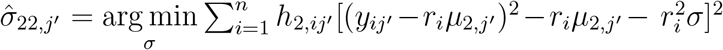, respectively. We set *w*_2,*ij’*_ = 1*/*Var(*ϵ*_2,*ij’*_) and 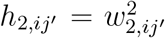, which either equates or approximates the inverse variance of the response variable to improve the statistical efficiency of the estimators [14]. As 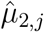 depends on true *µ*_2,*j’*_ and *σ*_2,*j*_*’j’* through *w*_2,*ij’*_ and 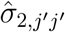 also depends on true *µ*_2,*j’*_, we propose an iterative procedure where we iterate between updating 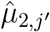 given 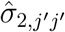 and vice versa. We propose to initiate the iteration with *w*_2,*ij’*_ = 1 (i.e. ordinary least squares estimator). The mean and variance for gene *j* can be estimated similarly from (2). A detailed algorithm of IRLS is included in Supplementary Algorithm 2.

Given these estimates, the second step of of scMultiMap estimates the covariance between peaks and genes with weighted least squares (WLS). Based on (4), we drive the regression equation

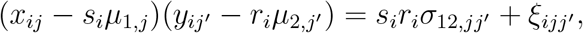

where mean-zero error terms *ξ*_*ijj’*_’s are independent across *i* for gene *j* and peak *j*’. We then propose to estimate *σ*_12,*jj’*_ via arg min 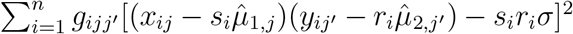. We set the weights *g*_*ijj’*_’s using *g*_*ijj’*_ =1^σ^*/*[Var(*x*_*ij*_)Var(*y*_*ij’*_)], which is the inverse variance of *ξ*_*ijj’*_ under the null hypothesis of independence between gene and peak, and we use weights *w*_1,*ij*_ and *w*_2,*ij’*_ from step 1 to calculate *g*_*ijj’*_.

For statistical inference on the association between peak and gene, we propose a test statistic based on the WLS estimator and analytically characterize its distribution under the null hypothesis. We define 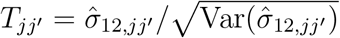, where 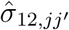’s are estimated with true *µ*_1*j*_, *µ*_2*j’*_’s. Under the null hypothesis of independence between gene expression and peak accessibility, we show that

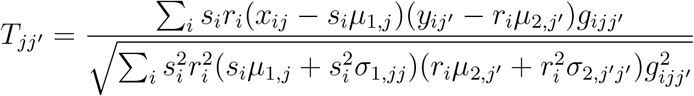

and that *T*_*jj’*_ follows a standard normal distribution (Supplementary Methods). This result facilitates the analytical calculation of test statistics and *p*-values without the need for computationally intensive sampling, and also ensures the theoretical validity of the test. In practice, we compute this test statistics by plugging in the IRLS estimates of mean and variance parameters from the first step, which are all consistent estimators.

When multiple subjects or biological samples are present in the single-cell multimodal data, we extend the model in (1) to model the variations in mean expression and accessibility across subjects. Suppose there are *K* subjects. For cell *i* = 1, …, *n* and subject *k* = 1, …, *K*, let 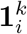 denote a binary indicator for subject *k*, i.e. 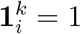 if cell *i* is from subject *k*, otherwise 0. We assume that

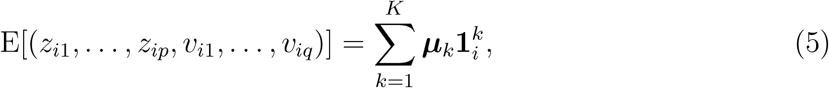

where *µ*_*k*_ denotes the mean for all genes and peaks in subject *k*. This implies the following moment condition for peak 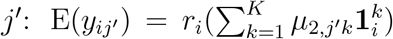, and similarly for gene *j*. We propose to estimate *µ*_2,*j’k*_’s and *µ*_1,*jk*_’s using a similar IRLS procedure and the estimation of variance and covariance follows analogously (Supplementary Methods).

### Other methods for statistical inference of peak-gene associations

We compared scMultiMap with two other methods: Signac and SCENT. For Signac, we used the peak-gene association method originally developed by [10], and implemented in the software Signac [9]. In benchmarking computational costs (Figure 1C), we used the *Link-Peaks* function in the R package Signac (v1.12.0) for the method Signac, and the R package SCENT (v1.0.0) for the method SCENT. SCENT was run without parallel computing to ensure a fair comparison with the other two methods.

In all other analyses, we used a re-implementation of *LinkPeaks()* to evaluate Signac. This re-implementation corrects a coding error in the *p*-value calculation in its original implementation and speeds up computation (Supplementary Methods). We applied SCENT with fixed numbers of bootstrap samples in numerical analyses (Supplementary Methods).

### Experiments for evaluating type I errors and power

To evaluate type I errors, we used a permutation-based procedure to generate null datasets where the gene expression and peak accessibility are independent. We used the gene counts as observed, and modified the observed peak counts following an approach that combines permutation and Poisson sampling as in [14]. This procedure maximally preserves the characteristics of real data, such as the mean accessibility of peaks, dependency of peak counts on sequencing depths, and the correlation of sequencing depths between modalities, to faithfully evaluate type I errors in real data. When multiple subjects are present in the same dataset (Figure 3), we applied the above procedure to cells from each subject separately, such that the variations in mean across subjects were preserved while any dependency between peaks and genes was removed. We obtained 100 independent replicates of null samples from this procedure and calculated the empirical *p*-values using the number of times the test statistic was rejected for all peak-gene pairs considered. To evaluate power, we simulated peak-gene pairs under model (1) with parameters estimated from real data, and incorporated model (5) for generating subject-specific means when multiple subjects are present in the dataset (Figure 3). We then evaluated the precision-recall curve by varying the cutoff of *p*-values for calculating false discovery rate (FDR) and power. More details can be found in Supplementary Methods.

### Reproducibility analysis

We performed two sets of reproducibility analysis between independent single-cell multimodal data, including on PBMC in Figure 2A and on brain samples in Figures 3D-E. For PBMC analysis, we considered [21] and [22] as biological replicates, which were generated with the same 10x instrument but with different biological samples. We also included a pair of technical replicates ([22] and [23]) that sequenced biological samples from the same individual but with different 10x instruments. We focused on CD14 monocytes as it is the most abundant cell types in these datasets. For brain analysis, we considered [32] and [33], which are two datasets independently generated by different labs, and evaluated the reproducibility on three most abundant brain cell types: excitatory neurons, inhibitory neurons and oligodendrocytes. In both analyses, two peak-gene pairs are defined as overlapped if two genes are the same and two peaks overlap in genomic ranges.

We performed two sets of consistency analysis with orthogonal data types, including on PBMC in Figure 2B and on brain samples in Figure 3C. For PBMC analysis, we combined three datasets [21, 22, 23] to maximize statistical power for detecting peak-gene pairs, and run scMultiMap and SCENT with indicators of datasets as covariates. In all analyses, we defined (candidate) cell-type-specific enhancer-gene pairs from orthogonal datasets following the original papers. In specific, for promoter capture Hi-C data [18], we used interactions with a CHiCAGO score *≥* 5 [62]; for H3K27ac HiChIP data [19], we used the significant interactions based on FitHiChIP [63] (FDR *<* 0.1); for cell-type-specific eQTL, we used cis-SNPs significant in Naive CD14 Monocytes (FDR *<* 0.05) [27]; for PLAC-seq, [3] profiles chromatin loops between promoters and distal regulatory regions in NEUN^+^ neuronal, OLIG2^+^ oligodendrocyte, PU.1^+^ microglia nuclei isolated from human brain tissues, and we used the provided data on enhancers, promoters, and interactomes on these cell types to define enhancer-gene pairs (Supplementary Methods). For datasets whose genomic locations are in hg19, we use *liftover* in R package rtracklayer (v1.62.0) to map the locations to hg38. In both reproducibility and consistency analysis, when multiple subjects are present in the data, we further adjusted Signac by its estimates on permuted null data that preserve across-subject variations. This is because correlations in Signac are known to be biased by mean [25, 14], and such systematic bias might cause artificially high reproducibility and consistency that are unfair to other methods with no systematic mean bias. Specifically, we computed Signac *p*-values using a background null distribution computed on permuted null data replicates, which corrects for the mean bias in Signac statistics.

### Analysis of cell-type-specific gene regulatory trios

Following the conceptual model in [28], we inferred cell-type-specific regulatory trios based on the associations between TF and target gene, TF and peak, peak and target gene using cells from the cell type of interest. We considered the highly expressed genes (mean expression ranks top 2,000) and highly accessible peaks (mean accessibility ranks top 20,000). We identified enriched motifs in highly accessible peaks using motifs from JASPAR 2020 database [64] and *FindMotifs* from R package Signac (v1.12.0). For each TF whose motifs are enriched and all highly expressed genes in the cell type, we constructed candidate trios using this TF, a highly expressed target gene, and highly accessible peaks located within the cis-region of the target gene (width 1Mb) and harbor the binding motifs of the TF. For all candidate trios, we then inferred the associations between TF and peak and peak and target gene with scMultiMap, and associations between TF and target gene with CS-CORE [14]. Based on the *p*-values, we prioritized trios whose all three edges are significant as cell-type-specific regulatory trios and evaluated the enrichment for biological processes among the target genes from all trios for each TF. More details can be found in Supplementary Materials.

### GO enrichment analysis

We used *gost* in R package gprofiler2 (v0.2.2) to perform enrichment analysis for GO biological processes with one-sided Fisher exact tests and selected driver GO terms with FDR*<* 0.05. We combined the selected terms manually based on the function of each cell type.

### Differential peak-gene associations

In differential analysis, we considered top 5,000 highly expressed genes and top 50,000 highly accessible peaks and the resulting candidate peak-gene pairs within the cis-region of width 1Mb in microglia and astrocytes. We applied scMultiMap while adjusting for the variations across subjects to cells from the control subjects and subjects with AD, respectively. To test the changes between two groups, we conducted a permutation analysis where we randomly permuted the disease label for 100 times and calculated *p*-values as the proportion of permutation replicates with the difference of covariance greater than that in observed data. In Figure 4B, we focus on gene-peak pairs that are significantly associated in either control or AD cells (raw *p*-value *<* 0.05) with significant difference between groups (raw *p*-value *<* 0.05) and the difference in correlation is greater than a magnitude of 0.2. We further intersected this set of genes with cell-type-specific differentially expressed genes from [32].

### LDSC analysis for heritability enrichment

AD heritability enrichment analysis was conducted using stratified linkage disequilibrium score (S-LDSC) regression [43] to determine if peaks from significant peak-gene pairs (raw *p*-value *<* 0.05) obtained by scMultiMap have higher heritability enrichment for AD in microglia compared to Signac and SCENT. S-LDSC calculates heritability enrichment in a stratum using the ratio of the proportion of heritability explained versus the proportion of SNPs in the stratum, based on GWAS summary statistics and an ancestry-specific reference panel. We used three AD GWAS summary statistics [41, 44, 45] and the European samples from 1000 Genomes data [65] as a reference panel. We estimated and tested the heritability enrichment of significant peaks from each method among all peaks considered in peak-gene association analysis, while adjusting for 97 functional annotations in baseline-LD v2.2. to best detect cell-type-specific heritability enrichment [66, 67].

### Preprocessing of fragment counts

We used fragment counts to quantify chromatin accessibility, which are more appropriate for count based modeling and can yield improved performance in downstream analysis compared to read counts [12, 13]. For three single-cell multimodal datasets on PBMC [21, 22, 23] and the dataset on postmorteum brain tissues [32], we called peak in each cell types respectively with MACS2 (bioconda v2.2.9.1) to obtain fragment counts and to maximize the discovery of cell-type-specific peaks. For the validation brain data [33], we calculated fragment counts based on read counts following the procedure in [13].

## Data Availability

All data used in this work are publicly available, including three datasets on PBMC from 10x Genomics [21, 22, 23] and two datasets on brain: GSE214979 from [32] and https://personal.broadinstitute.org/bjames/AD_snATAC/MFC/ from [33]. The detailed summary of each data set is in Supplementary Table 1.

## Code Availability

The R package that implements scMultiMap is available on Github: https://github.com/ChangSuBiostats/scMultiMap.

## Acknowledgement

Su was supported in part by the National Center for Advancing Translational Sciences of the National Institutes of Health under Award number UL1TR002378. The content is solely the responsibility of the authors and does not necessarily represent the official views of the National Institutes of Health. Jin was supported, in part, by the National Institutes of Health (NS111602 and HD104458). Zhang was supported by NSF grants DMS 2210469 and 2329296.

## Author Contributions

C.S. and J.Z. designed research; C.S. and D.L. performed research and analyzed data; C.S. contributed analytic tools; P.J. provided feedback on real data analysis; C.S., D.L. and J.Z. wrote the paper; C.S. and J.Z. jointly supervised the work.

