## Supplementary Information for "Cell-type-specific mapping of enhancers and target genes from single-cell multimodal data"

Details of defining cell-type-specific enhancer-gene pairs from orthogonal datasets 7

**Supplementary Table** **7**

**Supplementary Figures** **8**

### Supplementary Methods

#### Details of scMultiMap method

Here, we derive the formula for $T_{jj'}$ in the main text based on its definition $T_{jj'} = \hat{\sigma}_{12,jj'}/\sqrt{\text{Var}(\hat{\sigma}_{12,jj'})}$. Let $\hat{\sigma}_{12,jj'}$ be the WLS estimate with true $\mu_{1j}, \mu_{2j'}$, we have

$$
\begin{aligned}
T_{jj'} &= \frac{\hat{\sigma}_{12,jj'}}{\sqrt{\text{Var}(\hat{\sigma}_{12,jj'})}} \\
&= \frac{\sum_i s_i r_i (x_{ij} - s_i\mu_{1,j})(y_{ij'} - r_i\mu_{2,j'}) g_{ijj'} / \sum_i s_i^2 r_i^2 g_{ijj'}}{\sqrt{\text{Var}(\hat{\sigma}_{12,jj'})}} \quad \text{WLS} \\
&= \frac{\sum_i s_i r_i (x_{ij} - s_i\mu_{1,j})(y_{ij'} - r_i\mu_{2,j'}) g_{ijj'} / \sum_i s_i^2 r_i^2 g_{ijj'}}{\sqrt{\sum_i s_i^2 r_i^2 \text{Var}[(x_{ij} - s_i\mu_{1,j})(y_{ij'} - r_i\mu_{2,j'})] g_{ijj'}^2 / (\sum_i s_i^2 r_i^2 g_{ijj'})^2}} \quad \text{Independence across } i \\
&= \frac{\sum_i s_i r_i (x_{ij} - s_i\mu_{1,j})(y_{ij'} - r_i\mu_{2,j'}) g_{ijj'}}{\sqrt{\sum_i s_i^2 r_i^2 \text{Var}[(x_{ij} - s_i\mu_{1,j})(y_{ij'} - r_i\mu_{2,j'})] g_{ijj'}^2}} \\
&= \frac{\sum_i s_i r_i (x_{ij} - s_i\mu_{1,j})(y_{ij'} - r_i\mu_{2,j'}) g_{ijj'}}{\sqrt{\sum_i s_i^2 r_i^2 \text{Var}(x_{ij} - s_i\mu_{1,j}) \text{Var}(y_{ij'} - r_i\mu_{2,j'}) g_{ijj'}^2}} \quad \text{under H}_0 \\
&= \frac{\sum_i s_i r_i (x_{ij} - s_i\mu_{1,j})(y_{ij'} - r_i\mu_{2,j'}) g_{ijj'}}{\sqrt{\sum_i s_i^2 r_i^2 (s_i\mu_{1,j} + s_i^2\sigma_{1,jj})(r_i\mu_{2,j'} + r_i^2\sigma_{2,j'j'}) g_{ijj'}^2}} \quad \text{Equation (2) and (3),}
\end{aligned}
$$

which is exactly the formula shown in the main text. As $\hat{\sigma}_{12,jj'}$ is a weighted least squares estimate, $T_{jj'}$ converges in distribution to a standard normal distribution.

Next, We describe the algorithm of scMultiMap. Let $\odot$ denote the Hadamard product, and $\mathbf{1}_m$ denote a vector of 1's of length $m$. Algorithms 1-3 are for scMultiMap, and the IRLS and WLS step in scMultiMap, respectively. We write the algorithms in the general case where there could be $K$ biological samples in the dataset, with $K = 1$ (i.e. no multiple biological sample present) as a special case. Steps are written in matrix algebra whenever possible to resemble our efficient software implementation.

**Algorithm 1** scMultiMap

**Input:** Gene count matrix $X = (x_{ij})_{n\times p}$, peak count matrix $Y = (y_{ij})_{n\times q}$, binary matrix $E = (e_{ik})_{n\times K}$, where $e_{ik} = 1$ if cell $i$ is from biological sample $k$.

$\boldsymbol{s} \leftarrow X\mathbf{1}_p, \boldsymbol{r} \leftarrow Y\mathbf{1}_q$.

// Step 1:

// Estimate mean and variance parameters for genes.

$\{\hat{\boldsymbol{\mu}}_{1,j}, \hat{\sigma}_{1,jj}\}_{j=1}^{p}, \{\boldsymbol{w}_{1,j}\}_{j=1}^{p} \leftarrow$scMultiMap_IRLS$(X, \boldsymbol{s}, E)$

// Estimate mean and variance parameters for peaks

$\{\hat{\boldsymbol{\mu}}_{2,j'}, \hat{\sigma}_{2,j'j'}\}_{j'=1}^{q}, \{\boldsymbol{w}_{2,j'}\}_{j'=1}^{q} \leftarrow$ scMultiMap_ IRLS$(Y, \boldsymbol{r}, E)$

// Step 2:

// Estimate peak-gene association and compute test statistics

$\{\hat{\sigma}_{12,jj'}, \hat{T}_{jj'}\} \leftarrow$ scMultiMap_WLS$(X, \boldsymbol{s}, \{\hat{\boldsymbol{\mu}}_{1,j}, \hat{\sigma}_{1,jj}\}_{j=1}^{p}, \{\boldsymbol{w}_{1,j}\}_{j=1}^{p}, Y, \boldsymbol{r}, \{\hat{\boldsymbol{\mu}}_{2,j'}, \hat{\sigma}_{2,j'j'}\}_{j'=1}^{q}, \{\boldsymbol{w}_{2,j'}\}_{j'=1}^{q})$

**Output:** $\hat{\sigma}_{12,jj'}, \hat{T}_{jj'}$ for $j = 1, \ldots, p, j' = 1, \ldots, q$

**Algorithm 2** scMultiMap_IRLS

**Input:** Count matrix $X = (x_{ij})_{n \times p}$ for $n$ cells and $p$ features (genes or peaks), sequencing depths $\boldsymbol{s} = (s_1, \ldots, s_n)^\top$, binary matrix $E = (e_{ik})_{n \times K}$, where $e_{ik} = 1$ if cell $i$ is from biological sample $k$.
Set $\Delta^{(0)} = 1$, $t = 0$, $D = (\boldsymbol{s}\mathbf{1}_K^\top) \odot E$. Let $\boldsymbol{x}_j$ be the $j$-th column of matrix $X$.
// Define weighted least suqares operators for estimating $\boldsymbol{\mu}_j$ and $\sigma_{jj}$, respectively.
$f_{\boldsymbol{\mu}_j}(\boldsymbol{w}) \coloneqq (D^\top \text{diag}(\boldsymbol{w}) D)^{-1}(D^\top \text{diag}(\boldsymbol{w})\boldsymbol{x}_j)$
$f_{\sigma_{jj}}(\boldsymbol{\mu}, \boldsymbol{h}) \coloneqq [(\boldsymbol{s}^2)^\top \text{diag}(\boldsymbol{h})\boldsymbol{s}^2]^{-1} \left\{(\boldsymbol{s}^2)^\top \text{diag}(\boldsymbol{h})[(\boldsymbol{x}_j - D\boldsymbol{\mu})^2 - D\boldsymbol{\mu}]\right\}$
// Initialize with ordinary least squares
**for** $j = 1, \ldots, p$ **do**
    $\hat{\boldsymbol{\mu}}_j^{(t)} \leftarrow f_{\boldsymbol{\mu}_j}(\mathbf{1})$, $(\hat{\sigma}_{jj})^{(t)} \leftarrow f_{\sigma_{jj}}(\hat{\boldsymbol{\mu}}_j^{(t)}, \mathbf{1})$.
**end for**
// Iteratively reweighted least squares
**while** $\Delta^{(t)} \geq 0.05$ **do**
    $t \leftarrow t + 1$
    **for** $k = 1, \ldots, K$ **do**
        // Regularize $\theta_j$ estimates for weighting
        $\hat{\theta}_k^{(t)} \leftarrow \text{median}_j\{(\hat{\sigma}_{jj})^{(t-1)}/(\hat{\mu}_{jk}^{(t-1)})^2\}$
    **end for**
    // Update $\boldsymbol{\mu}_j, \sigma_{jj}$ estimates
    **for** $j = 1, \ldots, p$ **do**
        $\boldsymbol{w}_j^{(t)} \leftarrow 1/[D\hat{\boldsymbol{\mu}}_j^{(t-1)} + (D\hat{\boldsymbol{\mu}}_j^{(t-1)})^2 \times \hat{\boldsymbol{\theta}}^{(t)}]$
        $\hat{\boldsymbol{\mu}}_j^{(t)} \leftarrow f_{\boldsymbol{\mu}_j}(\boldsymbol{w}_j^{(t)})$
        $\boldsymbol{h}_j^{(t)} \leftarrow 1/[D\hat{\boldsymbol{\mu}}_j^{(t)} + (D\hat{\boldsymbol{\mu}}_j^{(t)})^2 \times \hat{\boldsymbol{\theta}}^{(t)}]^2$
        $(\hat{\sigma}_{jj})^{(t)} \leftarrow f_{\sigma_{jj}}(\hat{\boldsymbol{\mu}}_j^{(t)}, \boldsymbol{h}_j^{(t)})$
    **end for**
    // Assess convergence
    $\Delta^{(t)} \leftarrow \max_j |\log \hat{\sigma}_{jj}^{(t)} - \log \hat{\sigma}_{jj}^{(t-1)}|$
**end while**
$\hat{\boldsymbol{\mu}}_j \leftarrow \hat{\boldsymbol{\mu}}_j^{(t)}, \hat{\sigma}_{jj} \leftarrow \hat{\sigma}_{jj}^{(t)}, \boldsymbol{w}_j \leftarrow \boldsymbol{w}_j^{(t)}$, for $j = 1, \ldots, p$
**Output:** $\{\hat{\boldsymbol{\mu}}_j, \hat{\sigma}_{jj}\}_{j=1}^p, \{\boldsymbol{w}_j\}_{j=1}^p$

**Algorithm 3** scMultiMap_WLS

**Input:** $X, \boldsymbol{s}, \{\boldsymbol{\mu}_{1,j}, \sigma_{1,jj}\}_{j=1}^{p}, \{\boldsymbol{w}_{1,j}\}_{j=1}^{p}, Y, \boldsymbol{r}, \{\boldsymbol{\mu}_{2,j'}, \sigma_{2,j'j'}\}_{j'=1}^{q}, \{\boldsymbol{w}_{2,j'}\}_{j'=1}^{q}$
Set $D = (\boldsymbol{s}\mathbf{1}_K^\top) \odot E$. Let $\boldsymbol{x}_j$ be the $j$-th column of matrix $X$.
**for** $j = 1, \ldots, p,\ j' = 1, \ldots, q$ **do**
  // Compute regression weights
  $\boldsymbol{g}_{jj'} = \boldsymbol{w}_{1,j} \odot \boldsymbol{w}_{2,j'}$, let $G = \mathrm{diag}(\boldsymbol{g}_{jj'})$
  // WLS estimator for covariance
  $\hat{\sigma}_{12,jj'} = \left[(\boldsymbol{s} \odot \boldsymbol{r})^\top G (\boldsymbol{s} \odot \boldsymbol{r})\right]^{-1} \left\{(\boldsymbol{s} \odot \boldsymbol{r})^\top G \left[(\boldsymbol{x}_j - D\boldsymbol{\mu}_{1,j}) \odot (\boldsymbol{y}_{j'} - D\boldsymbol{\mu}_{2,j'})\right]\right\}$
  // Compute test statistics.
  $$\hat{T}_{jj'} = \frac{(\boldsymbol{s} \odot \boldsymbol{r})^\top G \left[(\boldsymbol{x}_j - D\boldsymbol{\mu}_{1,j}) \odot (\boldsymbol{y}_{j'} - D\boldsymbol{\mu}_{2,j'})\right]}{\sqrt{(\boldsymbol{s} \odot \boldsymbol{r})^\top \mathrm{diag}\left[(D\boldsymbol{\mu}_{1,j} + \boldsymbol{s}^2\sigma_{1,jj}) \odot (D\boldsymbol{\mu}_{2,j'} + \boldsymbol{r}^2\sigma_{2,j'j'}) \odot \boldsymbol{g}_{jj'}^2\right](\boldsymbol{s} \odot \boldsymbol{r})}}$$
**end for**
**Output:** $\hat{\sigma}_{12,jj'}, \hat{T}_{jj'}$ for $j = 1, \ldots, p,\ j' = 1, \ldots, q$

#### Evidence for overdispersion and variations across biological samples

To demonstrate overdispersion in peak counts in addition to Poisson, we performed goodness of fit test against Poisson for peak counts following the analysis for gene counts in [1]. We first randomly sampled 2,000 highly accessible peaks with probability inverse proportional to the density of mean accessibility, such that the sampled peaks evenly cover the mean accessibility levels [2]. For each of the sampled peak, we fitted a Poisson GLM model with sequencing depths as an offset, calculated the randomized quantile residuals with *statmod::qresid* in R [3], and performed a goodness of fit test using the residuals. We then used $p$-values from the goodness of fit test as evidence that Poisson models are insufficient to describe peak counts, and variance of quantile residuals as descriptive statistics for quantifying overdispersion in peak counts. We performed this analysis for three single-cell multimodal datasets on PBMC from 10x Genomics: PBMC_10k [4], PBMC_10k_nextgem [5], PBMC_10k_nextgem_X [6] and four abundant cell types in each of the datasets. The results are shown in Supplementary Figure 2.

To demonstrate the variations across biological samples, we quantified the variance in underlying abundance explained by the set of sample indicators $\{\mathbf{1}\{\text{cell } i \in \text{sample } k\}\}_{k=1}^{K}$ for genes and peaks respectively using F-test statistics. We then tested if the variance explained is equal to 0 using bootstrap hypothesis testing. We calculated $p$-values with 1,000 bootstrap samples for the genes and peaks and 3 abundant brain cell types analyzed in Figure 3, and the results are shown in Supplementary Figure 3.

To demonstrate that the mean variations tend to be correlated between peaks and genes, we evaluated the $p$-values for the Pearson correlations of variations between modalities (i.e.

$\text{Cor}(\hat{\boldsymbol{\mu}}_{1,j}, \hat{\boldsymbol{\mu}}_{2,j'})$ for $\hat{\boldsymbol{\mu}}_{1,j}$, $\hat{\boldsymbol{\mu}}_{2,j'}$ as defined in Algorithm 1) using a two-sided test implemented by *cor.test* in R. Supplementary Figure 4 shows the distribution of these $p$-values for the peak-gene pairs in 3 abundant brain cell types analyzed in Figure 3.

#### Implementations of Signac and SCENT

At the time of the study, the *LinkPeaks* function implemented by Signac mistakenly computes the two-sided $p$-value to be half of the correct value (see `https://github.com/stuart-lab/signac/blob/8ecdde2/R/links.R#L481C29-L481C30`). As a result, the $p$-value generated by Signac is bounded between 0 and 0.5, yielding artificially inflated type I errors in peak-gene association inference. We re-implemented *LinkPeaks* to correct for this coding error. We also separated the computation of background null distribution from the calculation of $p$-values, such that the null distribution can be fixed for all data replicates in permutation and simulation experiments (Figures 1A-B), speeding up the computation. For SCENT, we fixed the number of bootstrap samples to be 500 in permutation and simulation analysis to avoid extreme computational costs, and 5,000 in real data analysis to obtain refined $p$-values for multiple testing adjustment.

#### Details of evaluating type I errors and power

We used data from CD14 monocytes in PBMC data [5] and data from oligodendrocytes in brain data [7] to conduct the numerical experiments in Figures 1A-B and 3A-B, respectively. For each cell type, we first selected the top 2,000 highly expressed genes and top 20,000 highly accessible peaks, and used a cis-region of width 1Mb to select candidate peak-gene pairs. To save computational costs, we randomly sampled 1,000 pairs when evaluating the type I errors and precision-recall curves. In simulation experiments, we applied scMultiMap to estimate the parameters $\{\mu_{1,j}, \mu_{2,j'}\}$ and $\{\sigma_{1,jj}, \sigma_{2,j'j'}\}$ for the mean and variance of underlying gene expression $z_{ij}$ and underlying peak accessibility $v_{ij'}$ respectively, and $\sigma_{12,jj'}$ for the covariance between gene $j$ and peak $j'$. We further set $\sigma_{12,jj'} = 0$ for those pairs whose $p$-values are greater than 0.05. When multiple subjects are present in the data, we specified heterogeneous $\mu_{1,jk}$'s across subjects $k = 1, \ldots, K$, such that the standard deviation across $K$ subjects is 20% of the average mean $1/K \sum_{k=1}^{k} \mu_{1,jk}$, and that the largest overdispersion $(\sigma^2_{1,jj}/\mu^2_{1,jk})$ is 1 across subjects. We repeated the same procedure for peaks. Given the parameters, we then used a Gamma-Poisson mixture model combined with a Gaussian copula [8] to simulate correlated counts for each peak-gene pair [9].

#### Details in the analysis of gene regulatory trios

To maximize statistical power for inferring gene regulatory trios in PBMC, we combined cells from three PBMC datasets from 10x Genomics [4, 5, 6] and run scMultiMap with binary indicators for datasets as covariates. We used the following significance thresholds on raw $p$-values of scMultiMap and CS-CORE [9] for prioritizing regulatory trios: 0.05 for CD14 monocytes in PBMC data and 0.001, 0.001, 0.001, 0.05 and 0.1 for excitatory neurons, inhibitory neurons, oligodendrocytes, astrocytes, and microglia in brain data. We considered the top 5 enriched GO:BP driver terms (BH-adjusted $p$-value $< 0.05$) for the top 10 TFs with the greatest number of enriched trios in each cell type. Terms with relevant biological functions of each cell type are highlighted in Figure 3F-G and Supplementary Figure 5.

#### Details of defining cell-type-specific enhancer-gene pairs from orthogonal datasets

We used the Supplementary Table 5 of [10] to extract enhancers, promoters and interactomes on neuronal cells, oligodendrocytes and microglia. We adapted the codes from [11] and defined a pair of interacting peaks that covers a cell-type-specific enhancer and a promoter to be a enhancer-gene pair in that cell type. We used the results on neuronal cells to validate the findings in excitatory and inhibitory neurons.

### Supplementary Table

Supplementary Table 1: Summary of single-cell (nucleus) multimodal data used in analyses.

| Data sets | 10X [4] | 10X [5] | 10X [6] | GSE214979 [7] | Brain data from Kellis and Tsai labs [11] |
|---|---|---|---|---|---|
| Tissue | PBMC | PBMC | PBMC | Brain | Brain |
| #cells/nuclei | 11,457 | 9,229 | 9,517 | 105,332 | 65,046 |
| #cell types | 16 | 14 | 15 | 8 | 8 |
| Median RNA depth | 3,810 | 2,833 | 2,664 | 9,593 | 3,097 |
| Median ATAC depth | 13,030 | 13,826 | 13,156 | 6,638 | 2,159 |
| #genes | 36,601 | 36,601 | 36,601 | 36,601 | 36,601 |
| #peaks | 155,060 | 140,253 | 140,460 | 380,154 | 258,671 |
| #samples | 1 | 1 | 1 | 15 | 19 |

### Supplementary Figures

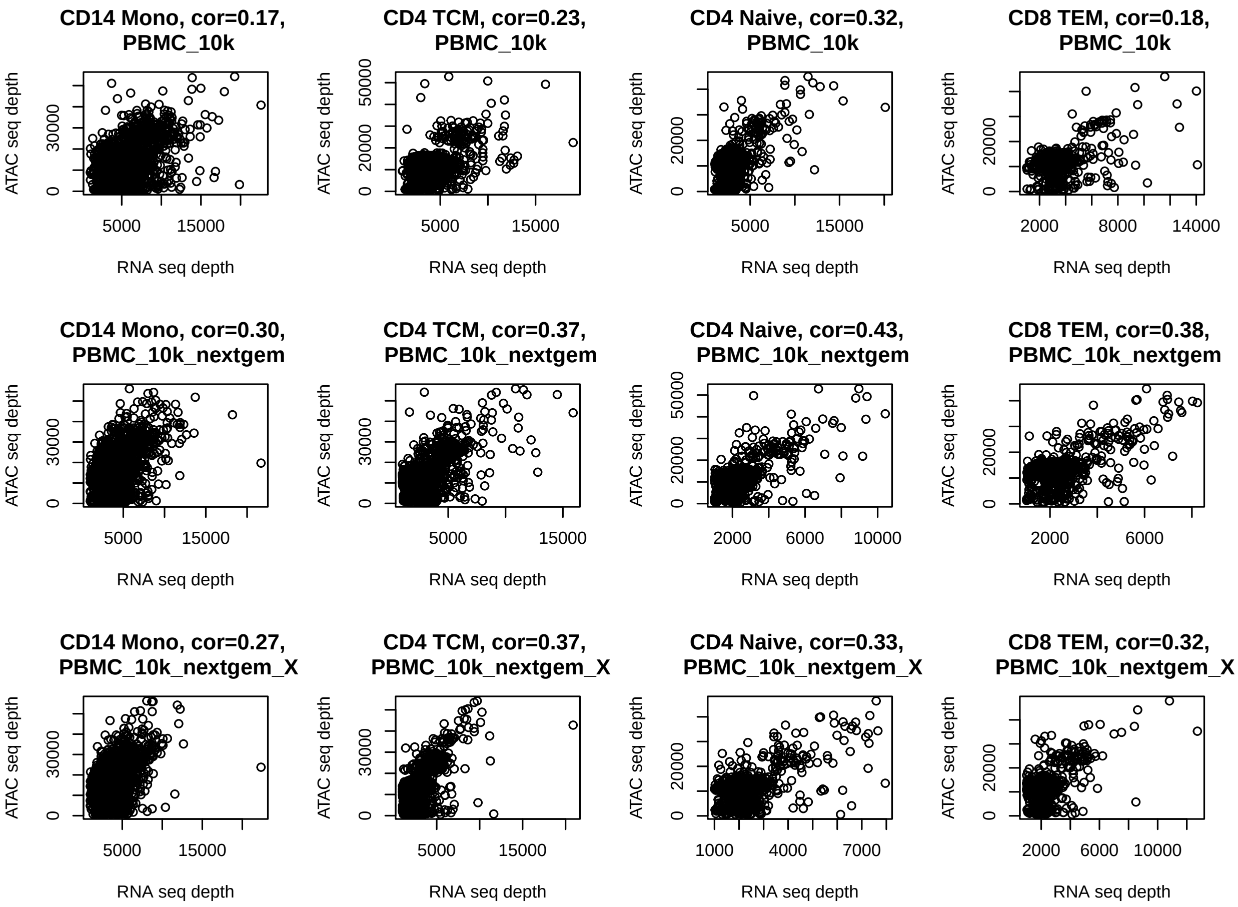

Supplementary Figure 1: Spearman correlations between sequencing depths of ATAC and RNA in three single-cell multimodal datasets on PBMC from 10x Genomics: PBMC_10k [4], PBMC_10k_nextgem [5], PBMC_10k_nextgem_X [6] across four abundant cell types in PBMC: CD14+ Monocyte (CD14 Mono), CD4+ Memory T Cell (CD4 TCM), CD4+ Naive T cell (CD4 Naive), CD8+ Effector Memory T Cell (CD8 TEM). Sequencing depths are computed as the total number of gene counts for RNA and peak counts for ATAC.

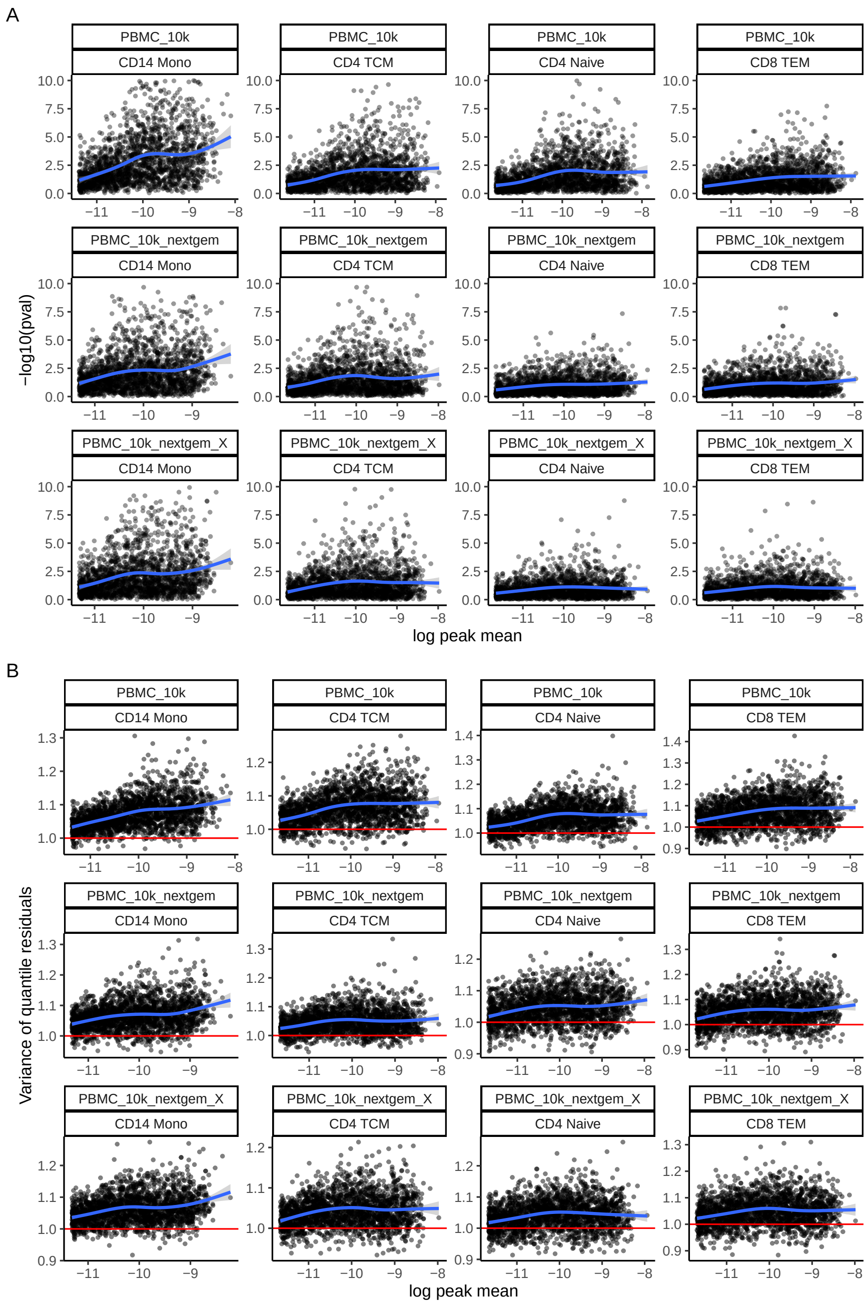

Supplementary Figure 2: Evidence for overdispersion in peak counts in three single-cell multimodal datasets from 10x Genomics and four cell types (same as Supplementary Figure 1). **A**. Results from the goodness of fit test against Poisson for peaks with varying mean accessibility. -$\log_{10} p$-values were truncated at 10 for visualization. **B**. Variance of quantile residuals for peaks with varying mean accessibility. Variance above 1 (denoted by the red line) indicates overdispersion, i.e. variance that cannot be explained by a Poisson model. Blue curves show the smooth fit by generalized additive models. See Supplementary Methods for more details on this analysis.

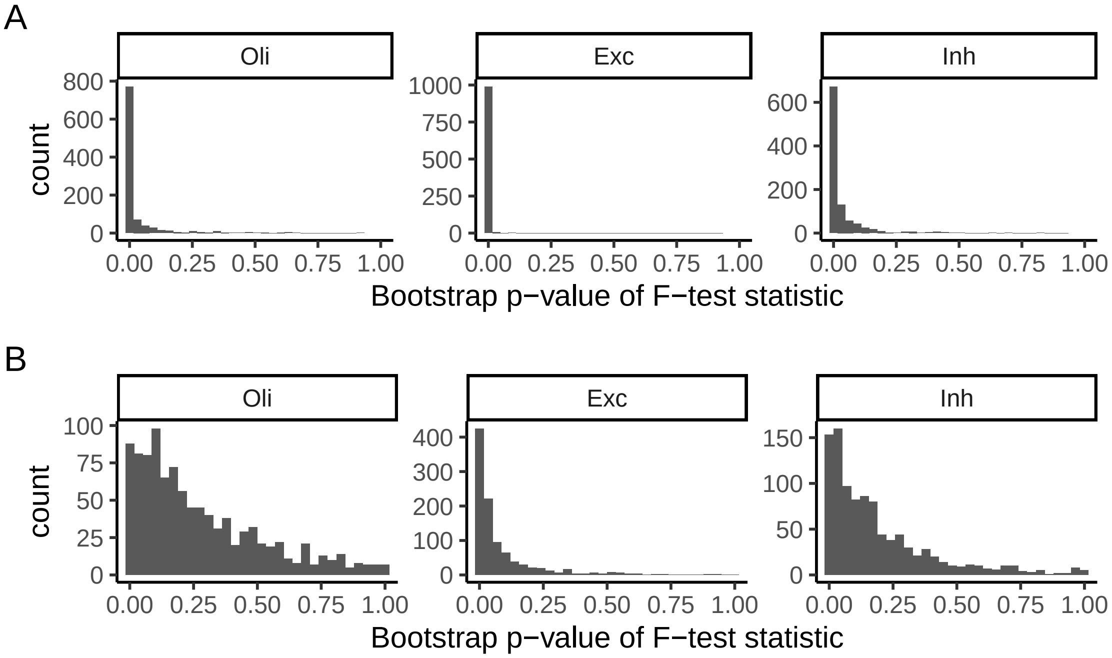

Supplementary Figure 3: Significant variations in mean gene expression (A) and mean peak accessibility (B) across biological samples based on single-cell multimodal dataset on brain from [7]. We evaluated the statistical significance of mean variations using one-sided bootstrap $p$-values of F-test statistics. See Supplementary Methods for more details on this analysis.

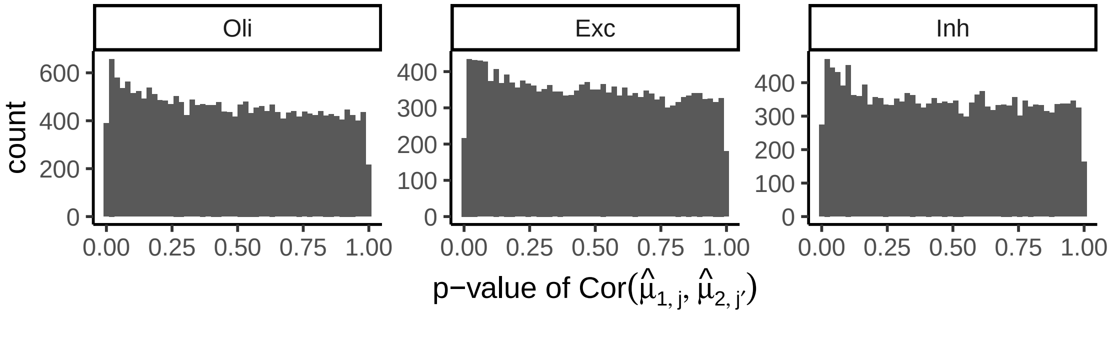

Supplementary Figure 4: Evidence for correlated variations between peaks and genes based on single-cell multimodal dataset on brain from [7]. We evaluated the statistical significance of correlations between peak and gene variations with two-sided $p$-values of Pearson correlations. See Supplementary Methods for more details on this analysis.

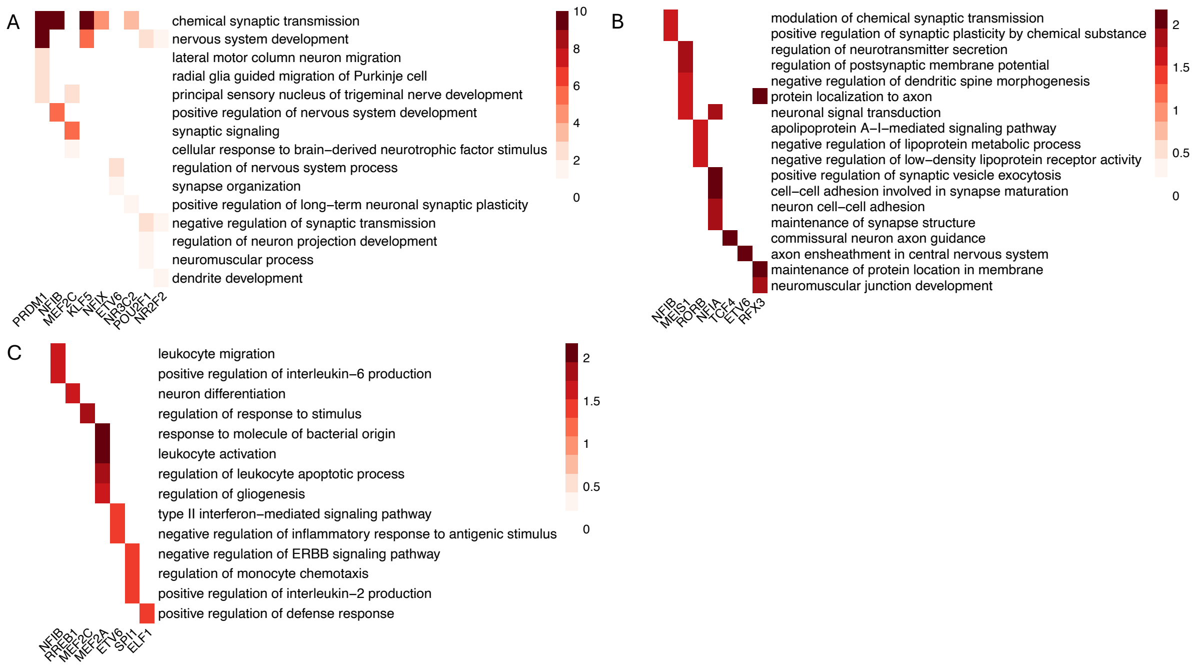

Supplementary Figure 5: Enrichment of GO biological processes among the target genes in trios identified for each TF in inhibitory neurons (**A**), oligodendrocytes (**B**), and microglia (**C**). Color intensity is given by BH-adjusted -$\log_{10}$ $p$-values from one-sided Fisher exact tests (values larger than 10 were set to 10).

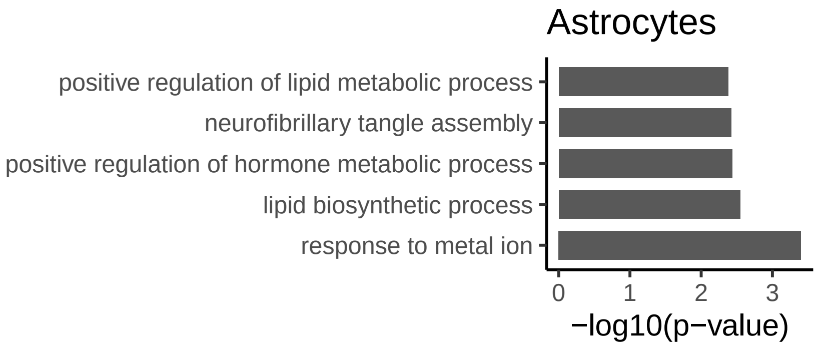

Supplementary Figure 6: Enrichment of GO biological processes among the genes from significantly differential peak-gene pairs in astrocytes (Supplementary Methods). BH-adjusted $p$-values from one-sided Fisher exact tests are shown.
